# Current state of bioacoustics research of vocal repertoires

**DOI:** 10.64898/2026.10.06.756971

**Authors:** Dori M. Grijseels, E. Pascal Malkemper

## Abstract

Knowledge of the unique call types produced by a species has historically been an important part of ecological research. However, the definition of a call type is not straightforward, and across species and studies different methods are employed to study call types and vocal repertories. Here, we analysed the methodology used to identify the vocal repertoires of 197 mammalian species across 220 studies. We found a large variability in the features extract from individual calls, methods to analyse spectrographic features, and methods to cluster call types. We further see a low uptake of novel methods for acoustic analysis and clustering. Lastly, we identify a lack of open data across the included studies, with less than 2% of studies providing the raw audio dataset underlying their paper. Based on these results, we call for increased standardization of methods and data transparency across the field.

## Introduction

Vocal repertoires, the variety of acoustic signals produced by animals, are of great importance to ecological and evolutionary research (Kershenbaum, Freeberg, and Gammon 2015b). For most species vocalizations are crucial signals through which individuals can signal their state, intention or feelings towards conspecifics and other species. As such, describing species’ vocal repertoire and the behavioral contexts of calls has historically been an important part of ecological research. Vocal repertoire and acoustic differences between calls may also be employed to determine or refine phylogeny (Burton and Nietsch 2010; Cap et al. 2008; Konrad and Geissmann 2006; Monticelli and Alencar-Jr 2021). Knowledge of call types have numerous further applications: acoustic recordings may aid conservation research (Teixeira, Maron, and van Rensburg 2019), by using acoustic signals to identify and count species, for example in heavily forested areas (Röper et al. 2014), or night-active species (Zimmermann et al. 2000). Lastly, vocal behavior is closely linked to other types of behavior, as well as an important welfare indicator, for example in farm settings (Manteuffel, Puppe, and Schön 2004). It is therefore not surprising that studying vocal repertoires has long been of interest to scientists.

However, defining a vocal repertoire is not trivial (Kershenbaum, Freeberg, and Gammon 2015b). In general, a vocal repertoire is defined as all call types made by multiple individuals of a species, but delineation of distinct call types is not always straightforward, a problem which has plagued the field for decades (Brown et al. 1978; Kershenbaum, Freeberg, and Gammon 2015b; Sainburg, Thielk, and Gentner 2020). What makes a separate call type differs by species and field, and may even depend on the individual researcher (Sainburg, Thielk, and Gentner 2020). For example, three different reports of elephant rumble vocalisations reported different repertoires, despite studying the same species (Soltis 2010). In some species the vocal repertoire is considered graded, meaning calls fall on a spectral continuum, and calls cannot be classified into distinct types (e.g. Barbary macaques (Hammerschmidt and Fischer 1998), bonobos (Keenan et al. 2020), and piglets (Tallet et al. 2013)). Increasing methods for the analysis of graded vocabularies are published (Cusano, Noad, and Dunlop 2021; Goffinet et al. 2021; Keen et al. 2021; Sainburg, Thielk, and Gentner 2020; Wadewitz et al. 2015), for example in Etruscan shrews (Langehennig-Peristenidou, Felmy, and Scheumann 2024) or orangutans (Erb et al. 2024), utilizing alternative quantifications of vocal complexity instead of classical descriptions of call types forming of vocal repertoires. Despite this, researchers may still prefer to identify distinct call types, and exclude intermediate calls that do not fit the categories (Chabout et al. 2015; Girard-Buttoz et al. 2022; Gursky 2015).

A further complication for comparative research in this field, is differences in terminology (De Gregorio et al. 2022; Vishwajith et al. 2026). A call may be defined as a single- or multi-unit vocalization, usually taken to be an acoustic signal produced through laryngeal activity in the case of mammals. However, whether a sequence of units should be considered a single call or multiple calls may differ per species. This delineation between a multi-unit call, or multiple single-unit calls, may be a simple interunit time threshold, for example any interval of longer than 200ms was considered to delineate a new call in squirrel monkeys (Schott 1975), or less than 400ms to delineate a new phrase in mouse USV song (Holy and Guo 2005). Another study defined all sound produced in a single expiration as part of the same ‘call’, regardless of short pauses within the sound produced (Minami and Kawamichi 1992). In some cases, sequences of syllables, also called motifs, are considered individual ‘call types’ for the purpose of establishing a repertoire, as for example in the case of murine ultrasonic songs (Holy and Guo 2005; Kalcounis-Rueppell, Metheny, and Vonhof 2006). This is similar to conventions in songbird research (Elie and Theunissen 2016; Loning, Griffith, and Naguib 2024), where ‘song’ may be considered a single call type. Lastly, some include the behavioral context in defining call types, thus splitting acoustically identical vocalizations into different types (e.g. (Jansen, Cant, and Manser 2012; Vanden Hole et al. 2014)), while others define calls purely on spectral properties.

Lastly, some many species are able to acquire new vocalizations through the process of vocal learning, including cetaceans (Janik 2014), pinnipeds (Reichmuth and Casey 2014), elephants (Poole et al. 2005), songbirds (Mooney 2009) and humans (Doupe and Kuhl 1999). In addition, vocalizations may be ‘culturally’ determined, meaning they are specific to a certain group. For example, in cetaceans, the repertoire may be clan specific (Gero, Whitehead, and Rendell 2016), and cultural turn-over may cause the disappearance or appearance of certain call types (Cantor et al. 2016). As a result, the vocal repertoire of a species may depend on the group of animals that is sampled, and may change over time.

This biological variation, in concert with inconsistencies in definition and methodology, makes interspecies comparisons particularly challenging. Although many recent publications have sought out more objective ways to quantify and describe vocal repertoires (Cusano, Noad, and Dunlop 2021; Erb et al. 2024; Pougnault et al. 2022; Sainburg, Thielk, and Gentner 2020; Wadewitz et al. 2015), no clear overview of the current methodology used in the literature, and the variety therein, is available. Here, we aim to address this issue, by performing a scoping review of peer-reviewed literature on the in-air vocal repertoires and call types of mammalian species. We selected in-air vocalizations only, excluding cetaceans, as the physics of underwater communication is different, further complicating comparisons. We extracted details on vocal repertoires and the methods of acquiring these for 197 different mammalian species, from 12 different orders, 49 different families. Overall, we aim to not only encourage discussion of better ways to consolidate cross-species results, but also petition for increased open science methods to enable further benchmarking and comparison in the future.

## Materials and Methods

### Search Strategy

We searched the PubMed database for (vocal repertoire) OR (acoustic repertoire), and . After removal of 5 duplicates, the titles and abstracts of the remaining 1212 results were prescreened according to our inclusion criteria using Rayyan (Ouzzani et al. 2016). Since many studies establish or analyse the vocal repertoire as a means to study further aspects of behavior or vocal communication, we included studies that did not explicitly note vocal repertoires in the abstract, but studied call types or vocal communication of a species.

We only included white primary literature in English in our main dataset, but also maintained a list of reviews, which were scanned for further primary literature. Additionally, further references were extracted from the retrieved papers, especially since older studies or studies with poor keywords may not have been included in our primary search, but were invariably referenced. Each received went through a further full-text screening, and those (re-)establishing a vocal repertoire, or analysing the repertoire of in-air call types in the adults of mammalian species were retained (for full inclusion criteria see supplementary A). When studies referenced a pre-established repertoire, and no further analysis or re-establishment of this repertoire was performed, the referenced study was extracted and added to the pre-text screening phase, and the referring study was excluded from the dataset (n=8). Studies investigating partial repertoires (e.g. one sex only, or in specific behavioral contexts) were included if multiple call types were distinguished and analysed. However, studies looking at individual differences within a single call type were excluded (n=6).

91 studies remained after preselection from the primary dataset and 115 studies were further extracted from the reviews and references, resulting in a total of 220 studies (Figure 1).

**Figure 1.**
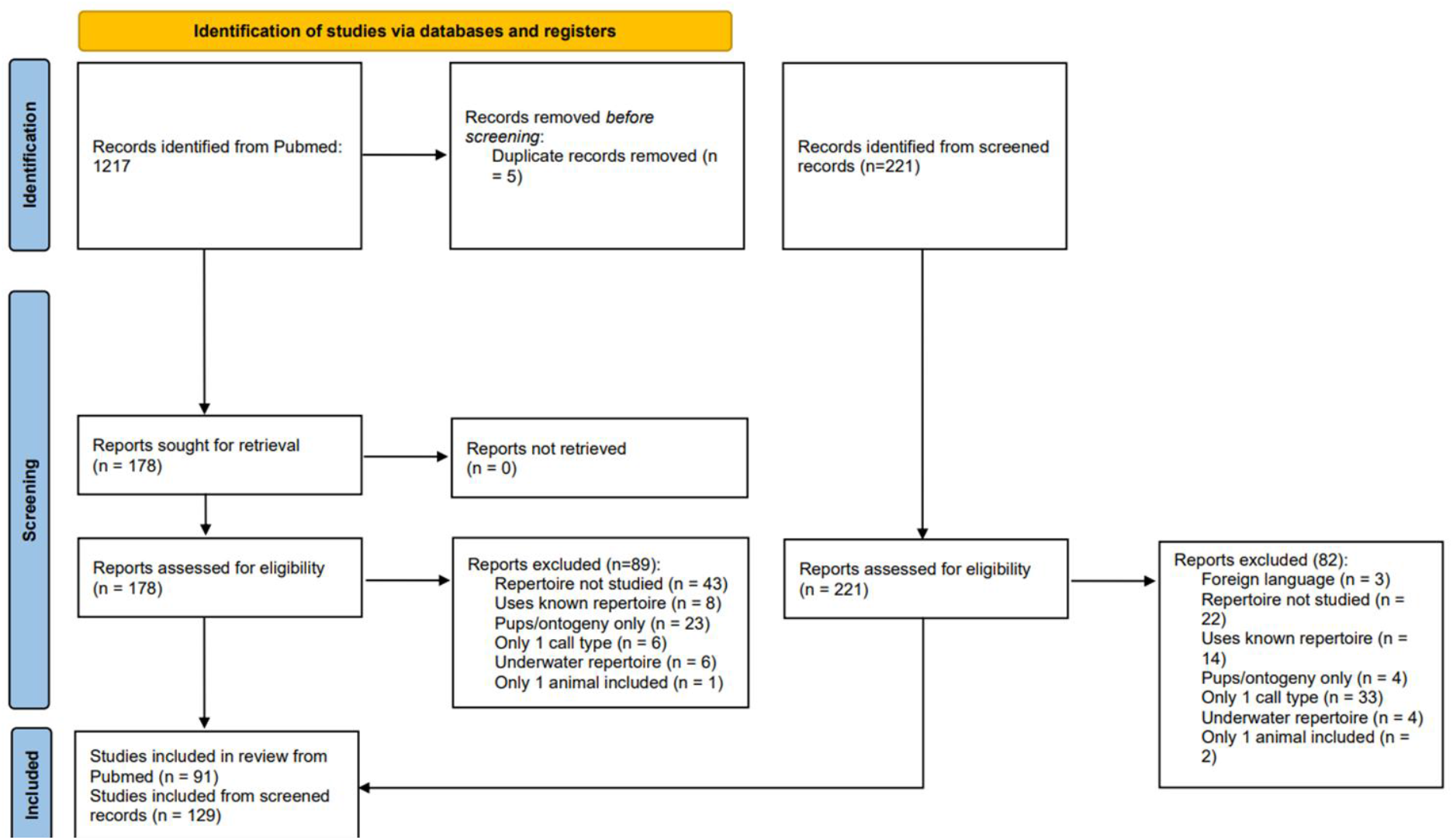
Prisma flow diagram for study inclusion.

### Information extraction

Metadata (such as publication year and journal) were extracted from the Rayyan database. Each of the 220 studies was manually examined for the relevant information about methodology. Preliminary processing was performed in Python using a custom script, figures were created in R v4.2.2 (R Core Team 2024). All data and code is available at https://github.com/DoriMG/vocal_repertoire_methods.

#### Study species

The reported species was recorded in English and Latin. Where updated nomenclature was available, this was used. For each species, the order and family it belonged to was further determined. If multiple species were analysed in the same paper, each species was noted separately.

#### Open access

For each report, it was recorded whether the report was available as open access. 2 records were made available post hoc through the Biodiversity Heritage Library, and are thus currently openly available, but were not considered open access.

#### Data sharing

The data sharing statement and available data were examined for each record. We further subdivided the availability into raw data, where the full dataset was freely available as raw audiofiles, preprocessed data, where the extracted features for each included vocalization were available, upon request, where the authors stated data was available upon request within the paper. Both cases where only partial data was available, or where only averages were given (i.e. not the information per vocalization) were not considered as not sharing data.

#### Recording location

The reported location of recording was determined. Where animals were trapped in the field and recorded in situ, this was considered as a field study. If animals were transported back to the lab and held here for recordings, these were included as lab recordings.

#### Context

When other animals were present during recording, these were considered group recordings. This included a wide range of experimental setups, including field recordings from wild groups, or pairing in the lab. Solitary context was defined as any recordings where conspecifics were not in visual or auditory range during the recording (either in the wild, or in the lab).

#### Number of animals

If the total number of recorded animals was reported, this was noted. In field recordings, the exact number of animals in the group was not always known, so no count was noted for these, even if an approximation was given. If multiple species were analysed in the same paper, the number for each species was noted separately.

#### Number of vocalizations

This represents the number of vocalizations included in the analysis. This number thus excludes recorded vocalizations that were excluded from analysis due to e.g. poor quality or unknown metadata.

#### Acoustic analysis software

The software or program used to extract features from acoustic vocalizations. In older papers, spectrograms were often produced using a spectrograph, and measurements were taken by hand, in which case the spectrograph was considered the analysis software. If multiple softwares or packages were used, all were included, thus each report could have more than one software.

#### Acoustic features

The number of individual unique features used in further analyses were determined from the records, when possible. When features were measured, but later excluded from analyses, e.g. due to high correlation between features, only the final number of included features were counted. The exact features were noted, and later sorted into larger categories.

#### Further analysis software

The software or program used to further analyse the vocalizations were noted. In cases where custom code was used, the programming language in which this was written was recorded. The first paper using LMA described the code as custom written, but formed the LMA package, and thus was included under this category. If multiple softwares or packages were used, all were included, thus each report could have more than one software.

### Terminology

As one of the issues with comparing studies is lack of consistent literature, we provide the definitions of terms as we will use them in this paper.

#### Vocal repertoire

All unique call types produced by the adults of a species.

#### Call

An acoustic emission, separated by considerable silence from other acoustic emissions. The length of the silence may depend on the species. A call may include multiple units (see below). Synonym: vocalization

#### Unit

An acoustic emission, separated by a short silence from other acoustic emissions, which together make up a single call. Synonym: syllable, note

#### Fundamental frequency

The lowest frequency component of an acoustic signal, in the case of tonal emissions.

#### Peak frequency

The frequency at which the highest amount of power is present in a frequency-power spectrogram. This may be the same as the fundamental frequency.

## Results

### Characteristics of included records

We extracted a total of 220 records, with publication years ranging from 1963-2025 (Figure 2a). Together these studies included 197 unique species, from 49 families spanning 12 orders. The largest number of studies included Primates and Rodentia (Figure 2b). When just considering the number of unique species studies per order, we see the same trend (S Figure 1a), with the highest number of species belonging to Primates and Rodentia orders. The studies were published in a wide range of journals, including species-specific journals (e.g. Primates), area-specific journals (e.g. Australian Mammalogy), bioacoustics journals (e.g. Journal of Acoustical Society of America), and general interest journals (e.g. PLOS One) (S Figure 1b).

**Figure 2.**
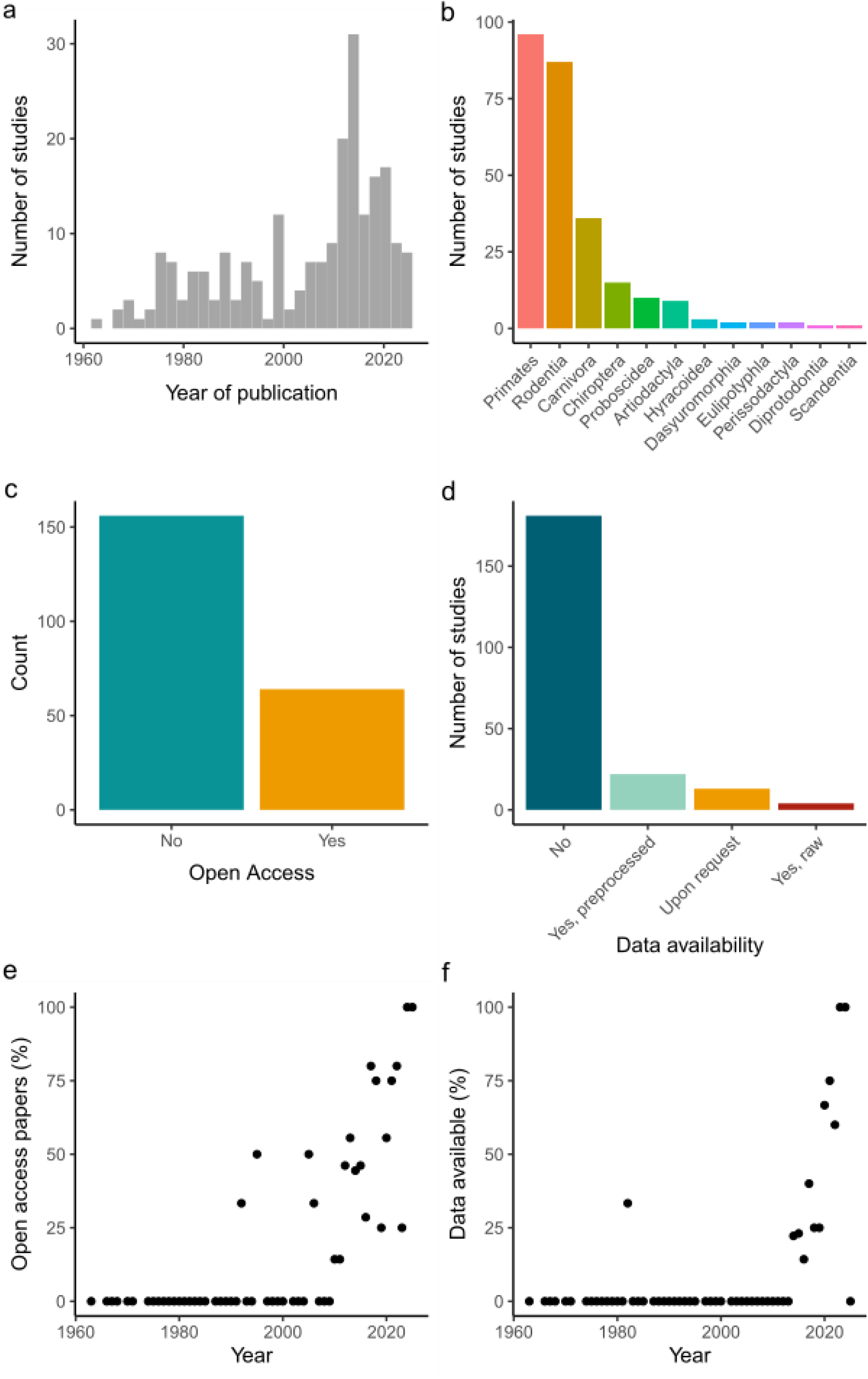
Characteristics of studies included in the review. (a) Histogram of the publication year of the studies. (b) The orders to which the studied species belonged to. (c) Open access status of the study. (d) The data availability for the study, divided into studies providing raw data, preprocessed data (e.g. extracted features per vocalization), or studies that indicated the raw data could be requested. (e) Trend in the open access status of papers as a function of the publication year. (f) Trend in the data availability of papers as a function of the publication year.

The minority (29.1%, 64 records, Figure 2c) of records were published open access, and even fewer studies shared data in any form (13.2%, 29 records, Figure 2d), with only 4 studies providing the full raw audio dataset freely online. However, we see an increase in open access, especially in recent years, with all 2024 (N=7) and the 2025 record having been published open access (Figure 2e). We see a similar upwards trend in the sharing of data, with all except for one study with any type of open data having been published after 2013 (Figure 2f).

### Methodology for collecting audio records

We next investigated what methods were used to collect audio recordings across our dataset. The majority of included records describe studies performed in the field (Figure 3a, 45.9%), with a large number of studies performed in labs as well (40%). Among the records were also studies using multiple recording locations, for example performing recording both in the field and in zoos (e.g. Mumm & Knörnschild, 2014). A small number of studies took place in other locations, such as farms or private homes. One study exclusively used published data, and was thus excluded from this analysis. Out of the 107 records that reported a social context, the majority of studies performed recordings in group contexts, meaning at least one conspecific was present at time of recording (Figure 3b, 157, 75.8%). Solitary recordings were often performed in combination with paired or grouped recordings (37, 17.8%), with a minority performing only solitary recordings (13, 6.3%). In solitary recording controlled stimuli were often used to elicit context-specific vocalizations, such as predators (Greene and Meagher 1998), or female urine (Hoffmann, Musolf, and Penn 2012). Across 140 reports for which the number of animals was reported, studies included a median of 22.5 individuals (range: 2-362, Figure 3c). Three studies with only a single research animal were excluded. We further recorded the number of vocalizations which were included in analyses, and found a median of 679 vocalizations (range: 20-583,237, Figure 3d). Although we do not see a clear increase in the number of animals studied over time (Figure 3e), there is a significant increase in the number of individual vocalizations used over time (Figure 3f).

**Figure 3.**
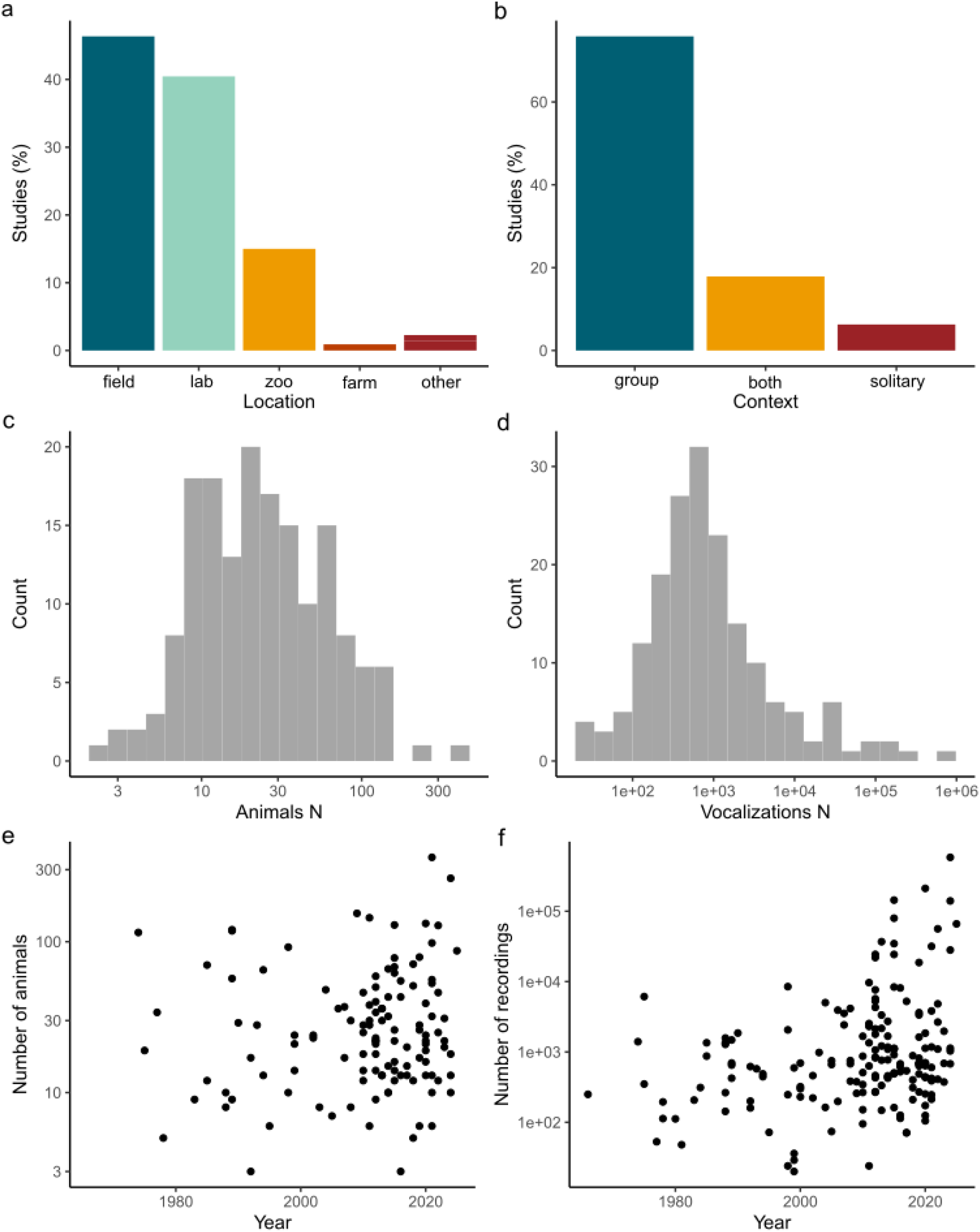
Recording methods across studies. (a) The location where recordings were performed, in some cases studies used more than one location. (b) Percentage of studies that recorded animals in presence of conspecifics (group), alone, or both. (c) Number of animals included in each study (median: 22.5, range: 2-362). (d) Number of vocalizations included in each study (median: 679, range: 20-583,23). (e) Number of animals included as a function of the publication date of the study. (f) Number of vocalizations included as a function of the publication year.

### Acoustic features used in vocal clustering

The number and types of features included in analyses can greatly impact the performance of clustering methods (Wadewitz et al. 2015). These features may either include acoustic features measured from the spectrogram, or latent features extracted using dimensionality reduction methods (Sainburg, Thielk, and Gentner 2020). For each study that performed further analyses on the repertoire, we determined what features (if any) were extracted from the recorded vocalizations to include in further processing.

99 (45.0%) studies performed acoustic analyses, of these 96 performed analyses based on acoustic features. Among these studies Avisoft-SASLab was the most common tool for acoustic feature extraction, followed closely by Praat (Boersma 2001) and Raven (Figure 4a). LMA was also used frequently (Schrader and Hammerschmidt 1997), though it should be noted that 80% of the papers using this package included the author of the package among the authors.

**Figure 4.**
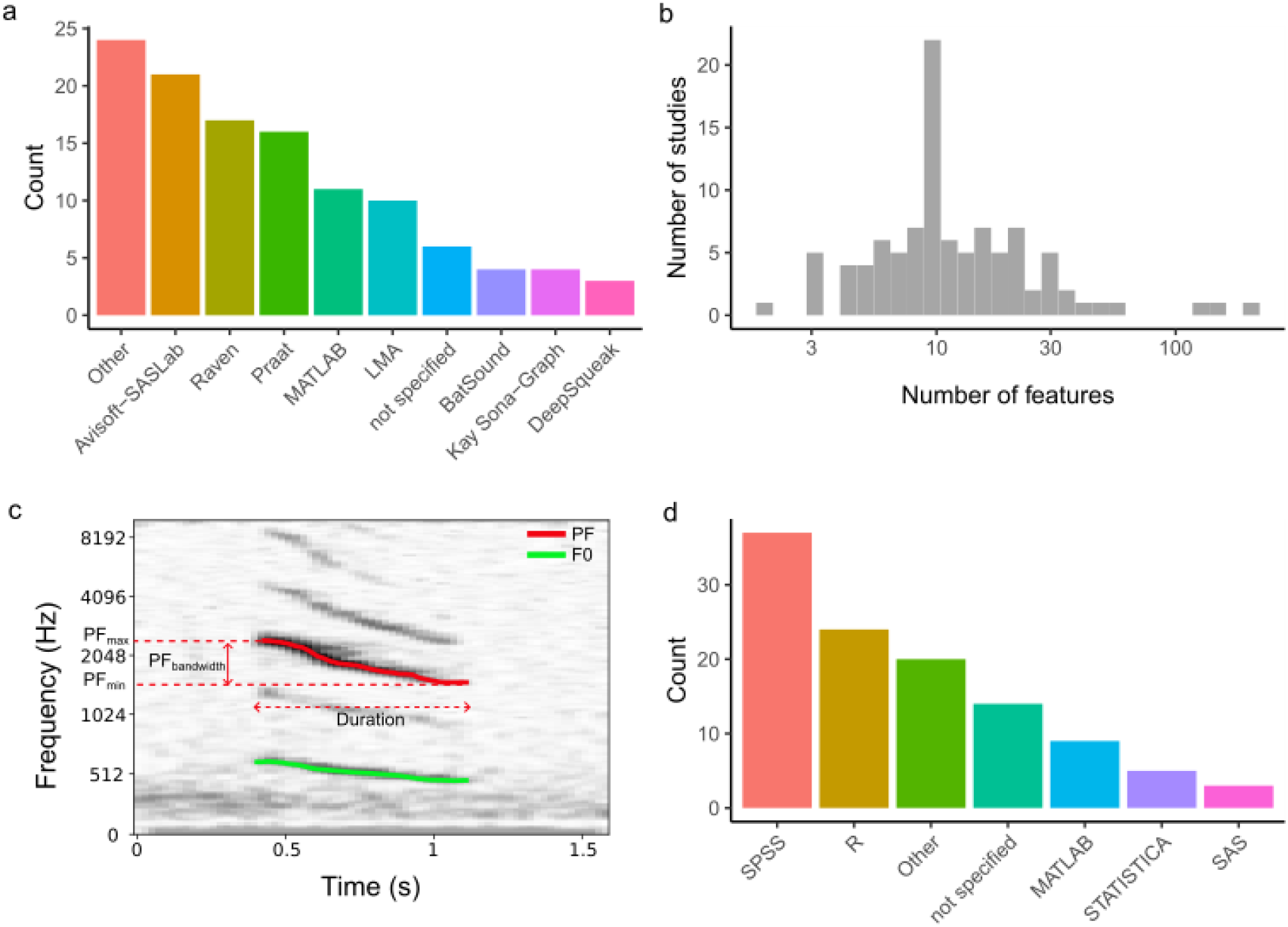
Analysis methods for bioacoustic data. (a) Softwares used for extracting features from vocalizations, quantified according to the number of studies that used a given software. (b) Number of features extracted for vocalizations from studies that performed further analyses on the vocalizations (median: 10, range: 2-217). (c) Example vocalization from an Ansell’s mole-rat (Fukomys anselli), showing peak frequency (PF, red) and fundamental frequency (F0) contours, as well as selected features that may be extracted from the PF contour. (d) Softwares used for further analyses of vocalizations, quantified according to the number of studies that used a given software.

A median of 10 acoustic features is used for further analysis (Figure 4b, range: 2-217) across the studies. We counted the occurrence of common features and feature classes (Table 1, see Figure 4c for example of feature measurements). Almost every study included duration of vocalizations (95.8%), with 34.4% including other temporal measures, such as time to maximum.

**Table 1.** Bioacoustic features extracted for further analysis.

| Feature type | Feature | Occurrence |
| --- | --- | --- |
| **Temporal** | Duration | 92 (95.8%) |
|  | Other (e.g. time to maximum frequency) | 33 (34.4%) |
| **Contour type (70 total studies)** | Fundamental frequency (F0) | 47 (67.1%) |
|  | Peak frequency (PF) | 21 (30.0%) |
|  | Unclear | 10 (10.4%) |
| **Spectrographic** | Minimum frequency | 62 (64.6%) |
|  | Maximum frequency | 67 (69.8%) |
|  | Start frequency | 35 (36.5%) |
|  | Center frequency | 19 (19.8%) |
|  | End frequency | 30 (31.3%) |
|  | Mean or median frequency | 36 (37.5%) |
|  | Peak frequency (frequency with the highest power) | 35 (36.5%) |
|  | Fundamental frequency | 13 (13.1%) |
|  | Bandwidth of frequency (usually max-min frequency) | 52 (54.2%) |
|  | Slope of frequency contour | 29 (30.2%) |
| **Multi-unit (48 total studies)** | Unit duration | 7 (14.6 %) |
|  | Number of units | 17 (35.4%) |
|  | Inter unit interval | 13 (27.1 %) |
|  | Unit rate (units/time) | 6 (12.5 %) |
| **Other** | Entropy | 13 (13.5%) |
|  | Amplitude | 12 (12.5%) |
|  | Power | 5 (5.2%) |
|  | HNR | 10 (10.4%) |
|  | Measures of harmonics (F1+ contour or mean, number of harmonics, etc) | 26 (27.1%) |

72.9% of studies identified a contour within the vocalization (see example Figure 4c). Of these studies, most used the F0 contour (67.1%), while 30.0% used a PF contour. In 10.4% of studies it was not specified whether the contour represented the F0 or PF contour. It should be noted that F0 and PF contour are often, but not always, the same for tonal vocalizations, but may differ drastically for atonal vocalizations, where F0 is not defined. Various spectrographic features were analysed, either from these contours or the overall frequency distribution, with the most common being maximum (69.8%) and minimum frequency (64.6%). The frequency at various points in the call were further included, in particular the start (36.5%), center (19.8%) and end (31.3%) frequency, as well as the mean or median frequency across the call (37.5%). The peak frequency (PF, 36.5%) is the frequency measured at the point with the highest power in the spectrogram, while the fundamental frequency (F0, 13.1%) is lower frequency in a harmonic tone. These two may be the same, but are not necessarily. Across studies including bandwidth (54.2%) we saw various ways of measuring this: in some cases the difference between maximum and minimum of the F0 or PF contour were used, whereas other studies measured a dark band in the overall spectrogram. Various slopes of the PF or F0 contour were measured (30.2%), usually slope across the entire call, or the start to max, and max to end slopes. Some studies (48, 50.0%) included multi-unit vocalizations, 14.6% of these measure the unit duration, 35.4% measure the number of units, 27.1% measure the inter unit interval, and 12.5% measure the unit rate. We further identified some common other measurements, such as measures of Wiener entropy (also called spectral flatness), either aggregate or average (13.5%), harmonics to noise ratio (HNR, 10.4%), and measure of non-fundamental harmonics, including the number of harmonics, and the frequencies or contours of these (27.1%). A small number also included amplitude measures (12.5%) or power measures (5.2%).

### Methods for quantifying vocal clustering

Lastly, we determined what analysis methods were used to cluster vocal types or analyse a proposed clustering. We only included studies here that used these methods specifically to analyse the vocal repertoire, though many studies used one or more of these methods for other purposes, such as classifying individuals, species, or treatments. Most studies used SPSS, followed by R (Figure 4d). By far the most-used method (Table 2) was some form of discriminant function analysis (DFA, 58.6%), which is typically used to quantify discrimination between the proposed call types. Various clustering methods (34.3%), including K-means clustering (14.1%), hierarchical clustering (8.1%) and other types of clustering (14.1%) such as Gaussian mixture models, were used for unsupervised clustering, with two studies employing multiple clustering methods. Multiple methods of dimensionality reduction were further employed (36.4%), including principal component analysis (PCA, 24.2%), multidimensional scaling (MDS, 9.1%), and other methods (7.1%). Only a minority of studies included here used any type of artificial neural network (ANNs, 6.1%).

**Table 2.** Analysis methods used for the study of vocal repertoires.

|  | Analysis method | Occurrence |
| --- | --- | --- |
| **Discrimination quantification** | DFA | 58 (58.6%) |
| **Clustering** | All | 34 (34.3%) |
|  | K-means or K-median | 14 (14.1%) |
|  | Hierarchical clustering | 8 (8.1%) |
|  | Other | 14 (14.1%) |
| **Dimensionality reduction** | All | 36 (36.4%) |
|  | PCA | 24 (24.2%) |
|  | MDS | 9 (9.1%) |
|  | Other | 7 (7.1%) |
| **Neural networks** | ANNs | 6 (6.1%) |

Studies using DFA usually report the results as the percentage of calls that were correctly classified, often but not always in conjunction with Wilk’s lambda. Across the 48 studies that reported accuracy of classification, an average accuracy of 81.7% was found (SFig 2A, range 55.3%-100%). The accuracy was not affected by the number of call types included in the analysis (SFig 2B).

## Discussion

In this review, we examined the methodology of vocal repertoire studies in a large variety of mammals. Although overall a minority of papers was published open-access and shared their data, we see a trend towards increased openness, with all 9 records from the 2 years having been published as open-access. Across studies and species there was no single method for defining and quantifying vocal repertories, as illustrated by the large variety of features used, and the variety in further analysis methods. However, we did find some throughlines, such as the inclusion of duration as a key feature, the use of various measures of fundamental or peak frequency, and the use of discriminant function analysis for repertoire clustering confirmation.

A minority of studies shared data in any form, with most data being only accessible preprocessed or upon request. As we saw such a variety in features extracted, this means it will be difficult to perform comparisons of vocalizations between studies, as it is very unlikely that the same features were extracted. Data being shared in a difficult to use manner is a common issue in ecology (Roche et al. 2015), and is not unique to bioacoustics. Although we further considered the 13 studies which indicated that data was available upon request as sharing their data, previous work has shown that this often does not actually lead to data availability (Tedersoo et al. 2021). Lack of data sharing is generally seen as an impediment to science overall (Tenopir et al. 2015), and we believe this to be the case in the field of bioacoustics as well. In recent years, several substantial bioacoustics datasets have been published (Best et al. 2025; Lamothe et al. 2025; Prat et al. 2017) which may serve as examples for standards in data sharing for bioacoustics. However, we also recognize barriers still exist (Reichman, Jones, and Schildhauer 2011), for example a lack of access to servers to store large amounts of bioacoustic data. Journals publishing the research (Koivisto and Mäntylä 2024) and the scientist performing it should work in concert to address these barriers, for example by engaging in targeted efforts to improve this infrastructure, as seen in other disciplines (Rübel et al. 2022)

Across the analysed report, we found a large variety in species and number of animals, number of vocalizations, recordings methods, features extracted and analysis methods. Each of these factors is likely to affect the identified vocal repertoire. The number of vocalizations recorded may impact which call types are observed. Particularly rare call types may happen to not occur in the dataset, causing an undercounting (Kershenbaum, Freeberg, and Gammon 2015a), though computational methods may counter this problem (Harris et al. 2016), Although we did not record details about the behavioral context of recording, this may additionally impact what vocalizations are observed. Certain calls, such as courtship calls or alarm calls, are only produced in particular contexts. As such, presenting of (fake) predators was common in studies on alarm calls (e.g. (Collier et al. 2020)), and presenting female urine was common in eliciting courtship vocalizations (e.g. (Hoffmann, Musolf, and Penn 2012)). Overall, obtaining a large number of vocalizations across individuals, ages, sexes, and (behavioral) context is likely the best way to increase the accuracy of a repertoire.

We found a large variety in the extracted features across the reports. While extracting the features it became clear that there was an inconsistency in naming across studies and tools. In reports it was not always clear what measurements were taken exactly. Across the studies that analysed the frequency contour, a substantial portion of the reports did not clarify whether the peak or fundamental frequency contour was extracted. Altogether, this causes significant issues for reproducibility. Standardized feature sets may aid in this, and they were occasionally used (Arnaud et al. 2023), but given the variety of vocal signals produced across species, it is difficult to find a single set that accurately describes all relevant aspects of all calls. In particular, atonal calls, calls that show non-linear phenomena, can often not be described using frequency contour (Muir et al. 2025), though some studies solve this by using different feature sets for tonal and non-tonal calls (Maust-Mohl, Soltis, and Reiss 2015). Some unique features, such as the number of frequency jumps in USV (Warren et al. 2021), do not apply to other call types or species at all, and thus should not be included in a universal feature set, even though they may be informative in certain use cases. Latent features, features extracted using machine learning techniques such as autoencoders, may be applied across all call types (Sainburg, Thielk, and Gentner 2020), but may not represent ecologically relevant characteristics of the calls, and so may not always be desirable for describing vocalizations. It is unlikely a standardized method for feature extraction can be used across all bioacoustic research, but improving descriptions and sharing code for the features used is likely to improve overall reproducibility.

In terms of further analysis methods, DFA was the most-used method to corroborate manual classifications, and a high accuracy in the classification was considered evidence for an accurate classification. However in studies that reported the accuracy per call type, the accuracy for some individual call types is very low, with less than 40% of calls being classified correctly for some call types (Bastos et al. 2015; Batist et al. 2023; Lisboa et al. 2021). Some studies examined misclassifications to determine similarities between certain classes (e.g. (Leger, Owings, and Gelfand 1980)). In addition, even when individual call types had a low accuracy, the classification was then rarely reconsidered based on the DFA results (but see (Eisinger, Scheibe, and Flaherty 2016)). Arnaud et al. (2023) compared DFA to other classification alternatives, and concluded these alternatives are often better and more accurate than DFA, particularly in Small, Unbalanced, Noisy, but Genuine (SUNG) datasets, which constitutes the majority of studies included here. In the reports included here, we did not see a wide uptake of the alternatives they tested, in particular Support Vector Machines (SVM), neural networks (NNs) and xgboost. It is clear that DFA has been a field standard for decades, but including better methods will likely benefit bioacoustic research going forward.

Bioacoustics research has seen incredible developments in recent years, with the increasing availability of machine learning techniques and computation power making analysing large amounts of data feasible. Several practical guides, with corresponding code, have additionally been published to help researchers make use of these techniques (Kershenbaum et al. 2025; Sainburg, Thielk, and Gentner 2020; Thomas et al. 2022), but so far we see a low uptake of these newer techniques. In the coming years, we hope increases in open-access publications, open datasets and uptake of new machine learning techniques will aid in significantly advancing the field of bioacoustics.

## Acknowledgements

We thank Ruth Scherger for aid in procuring selected records included in this study.

## Declaration of generative AI use

The authors report generative AI was not used in their research or preparation of this manuscript.

## Author Contributions

DMG – Conceptualization, Formal analysis, Investigation, Visualization, Writing – original draft, Writing – review & editing; EPM: Supervision, Writing – review & editing

## Supplementary information

**Figure S1.**
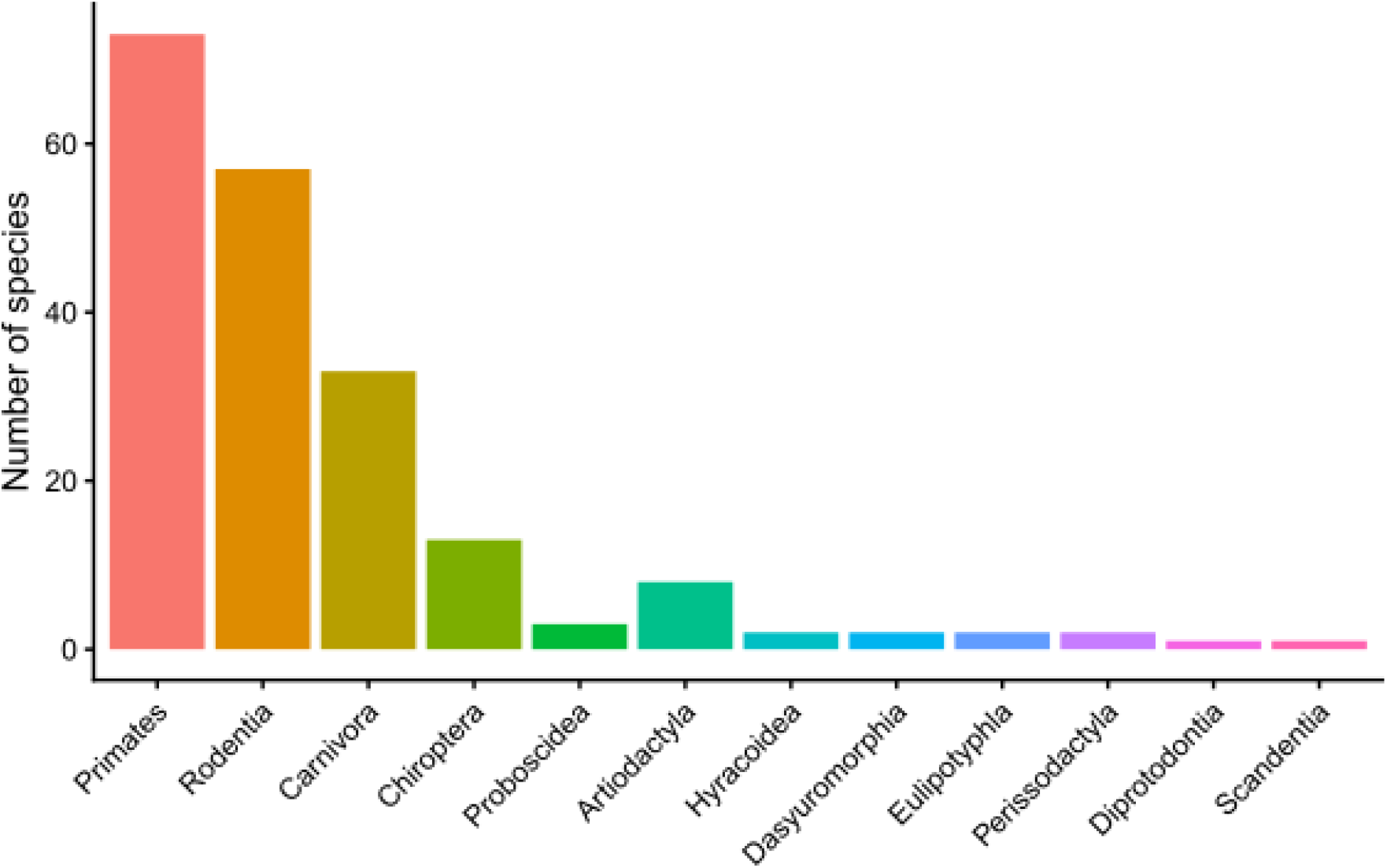
Study species. The mammalian orders to which the studied species belonged, counted by the number of unique species per order included in the dataset.

**Table S1.**
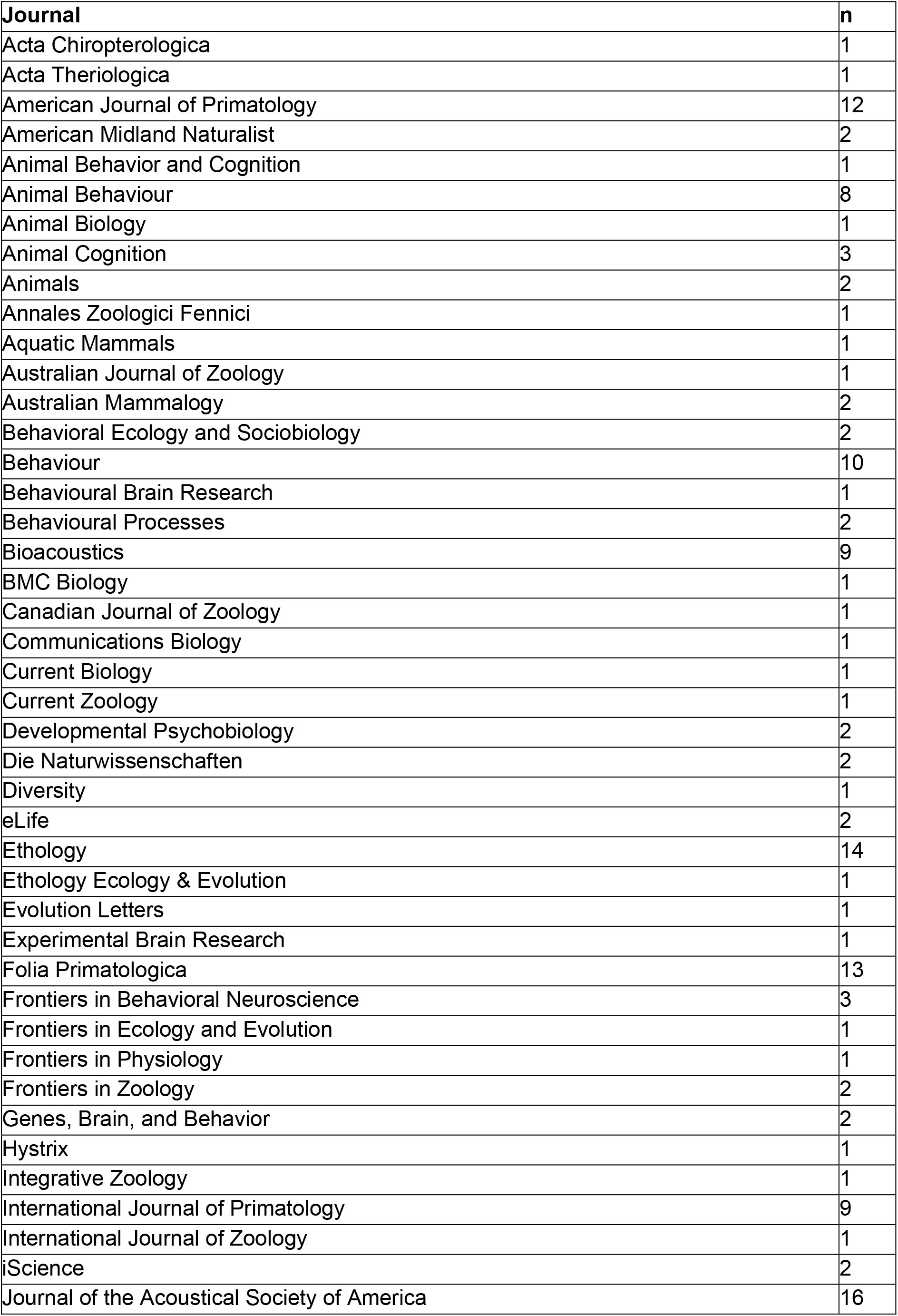
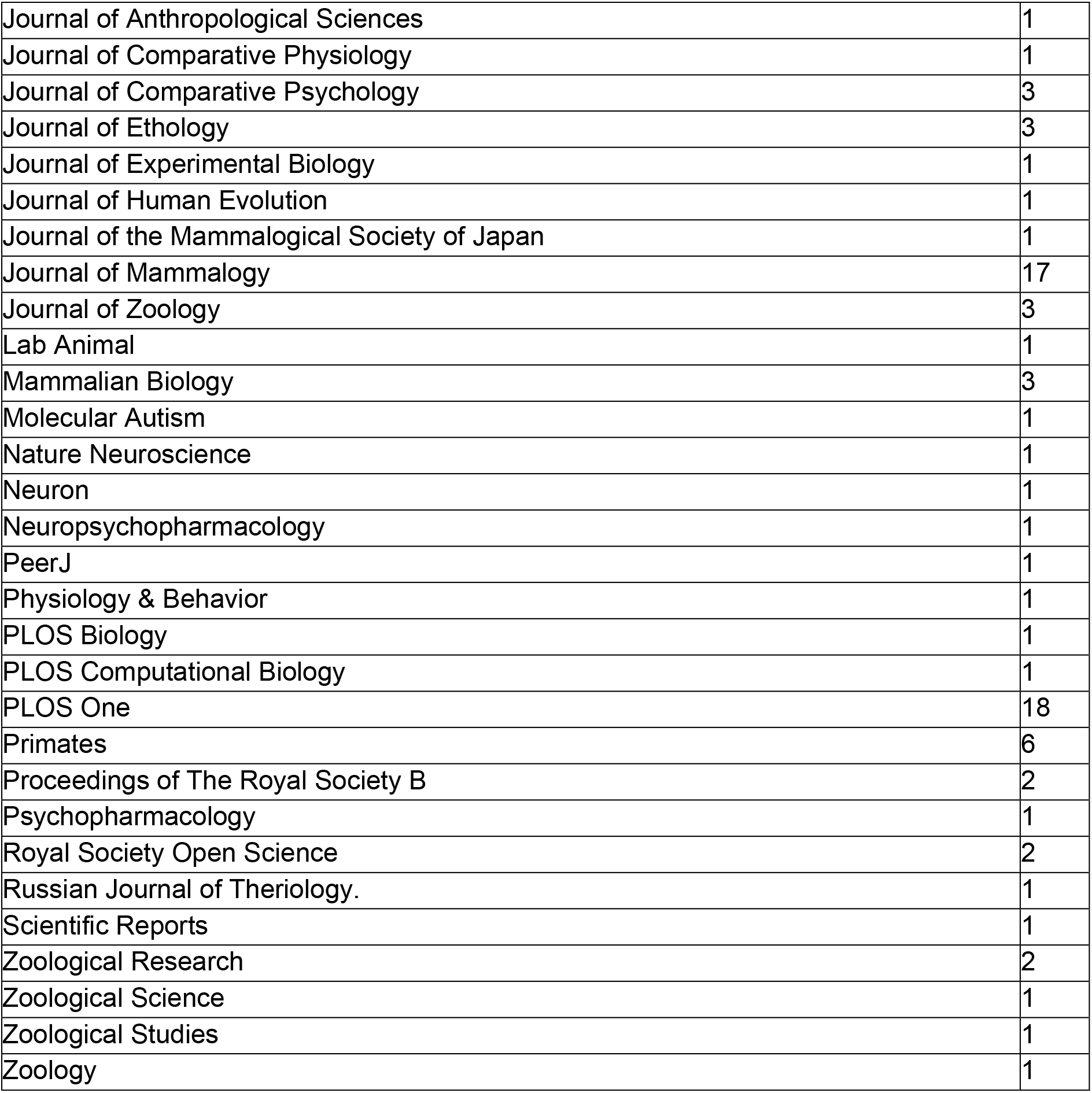
Publication outlet of the studies.

**Figure S2.**
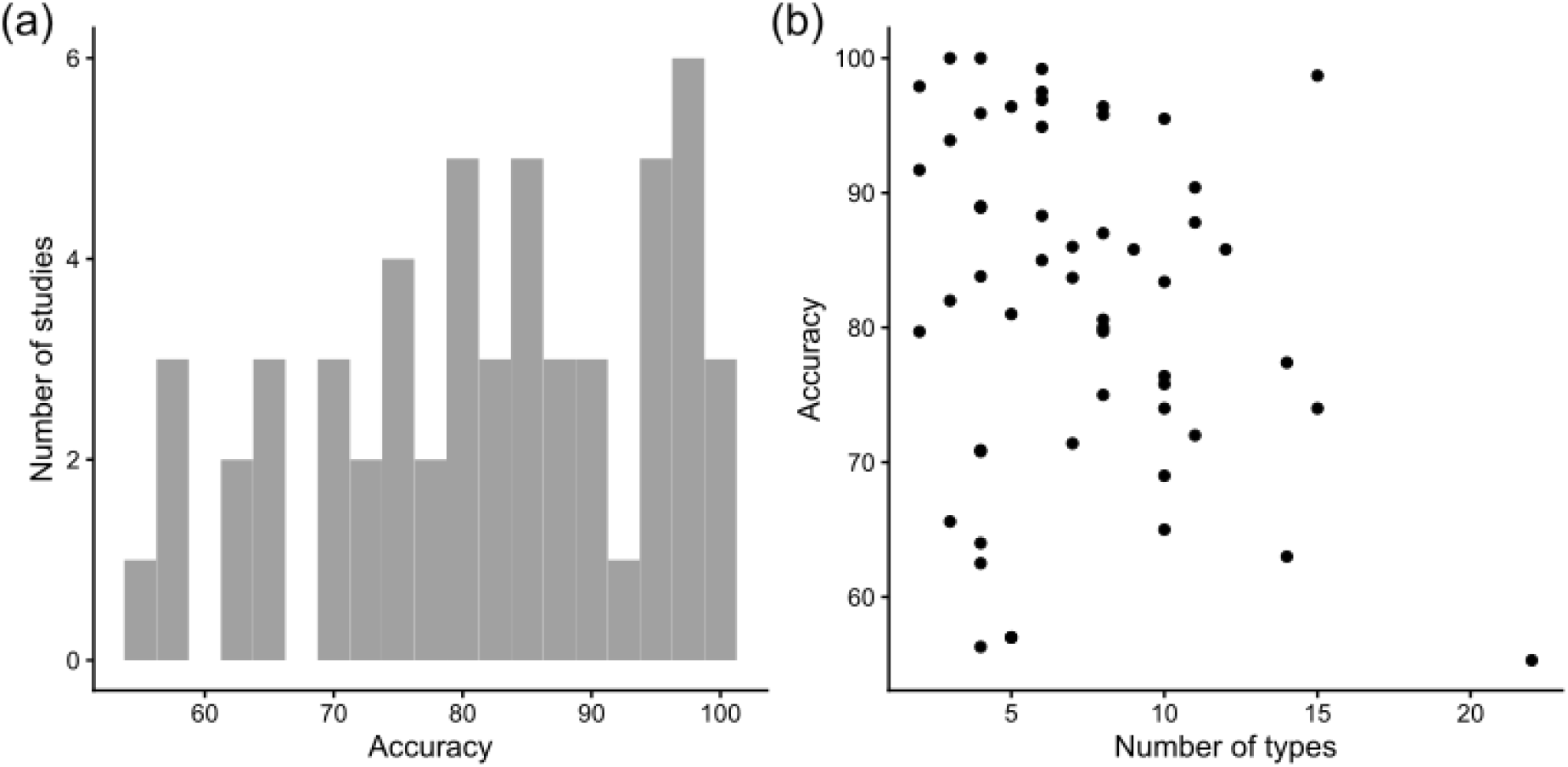
DFA accuracy performance. (a) The accuracy reported for DFA analyses. (b) The accuracy for DFA analyses as a function of the number of call types classified.

## Appendix A: Inclusion criteria

1. Record should be an English-language peer-reviewed primary research paper
2. Studies should include terrestrial vocalisations of mammalian species

a. Terrestrial vocalisation of semi-aquatic species (e.g. pinnipeds) are included
b. In-water vocalisations of aquatic or semi-aquatic species are excluded
3. Studies should include comparison of >1 vocalisation type

a. Studies including multiple types within a single context (e.g. courtship) are included
4. Studies should include >1 individual

a. Studies only including one sex are included, as long as multiple adult animals are included.
5. Studies should include adults of the species

a. Studies only including pups/juveniles are excluded
b. Studies of ontogeny, where no adults or species repertoire is established are excluded
6. Studies should perform acoustic or spectrographic assessment or analyses

a. Studies performing qualitative and/or subject assessment (e.g. visual inspection of spectrograms) are included
b. Studies purely analysing timing are excluded
7. Studies should establish or re-establish a separation of vocal types

a. Studies using a previously described repertoire are excluded, but the original established repertoire is considered for inclusion
b. If a previous repertoire is used, but is altered or adapted, both studies may be included.

## Appendix B: Scanned reviews

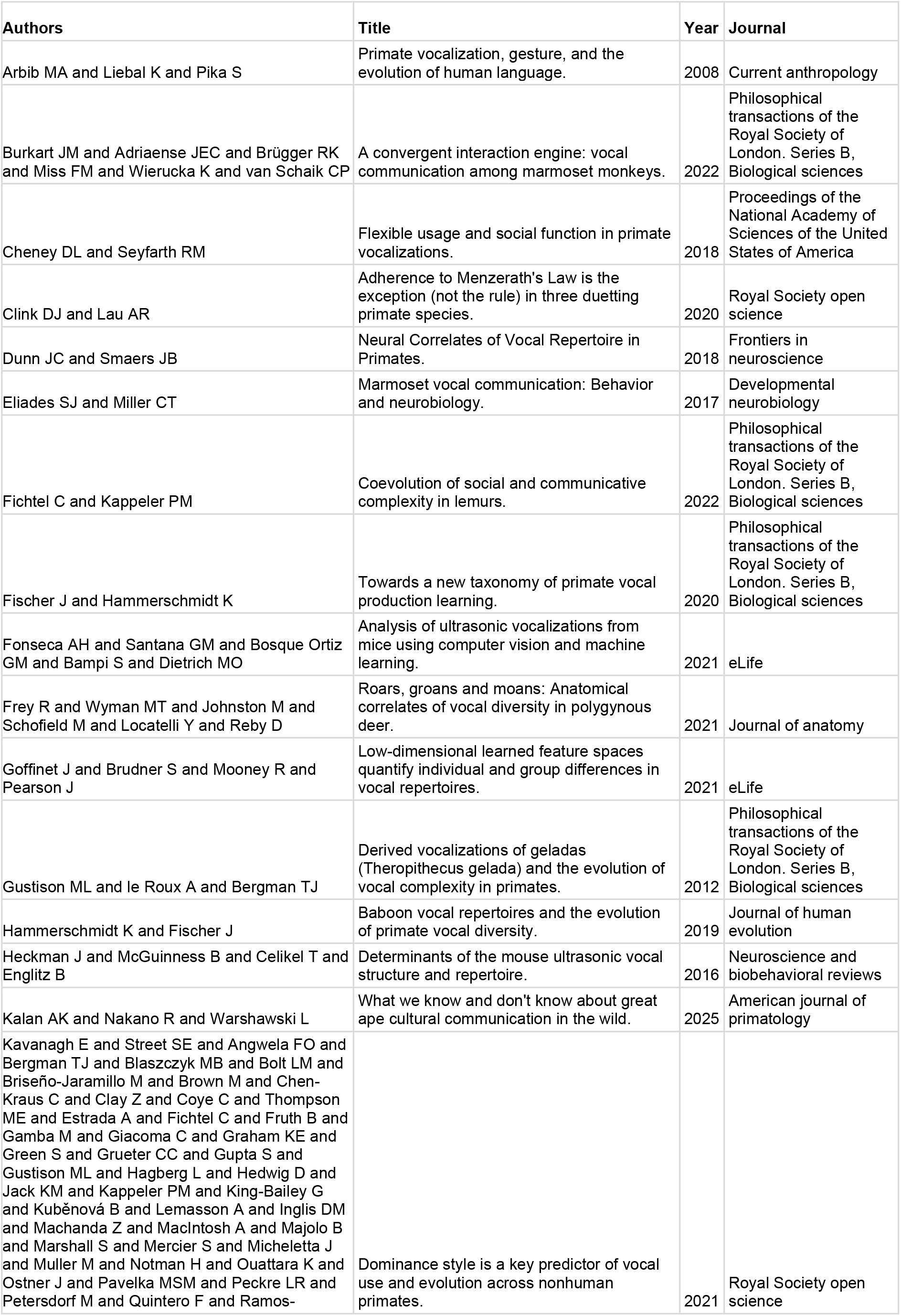
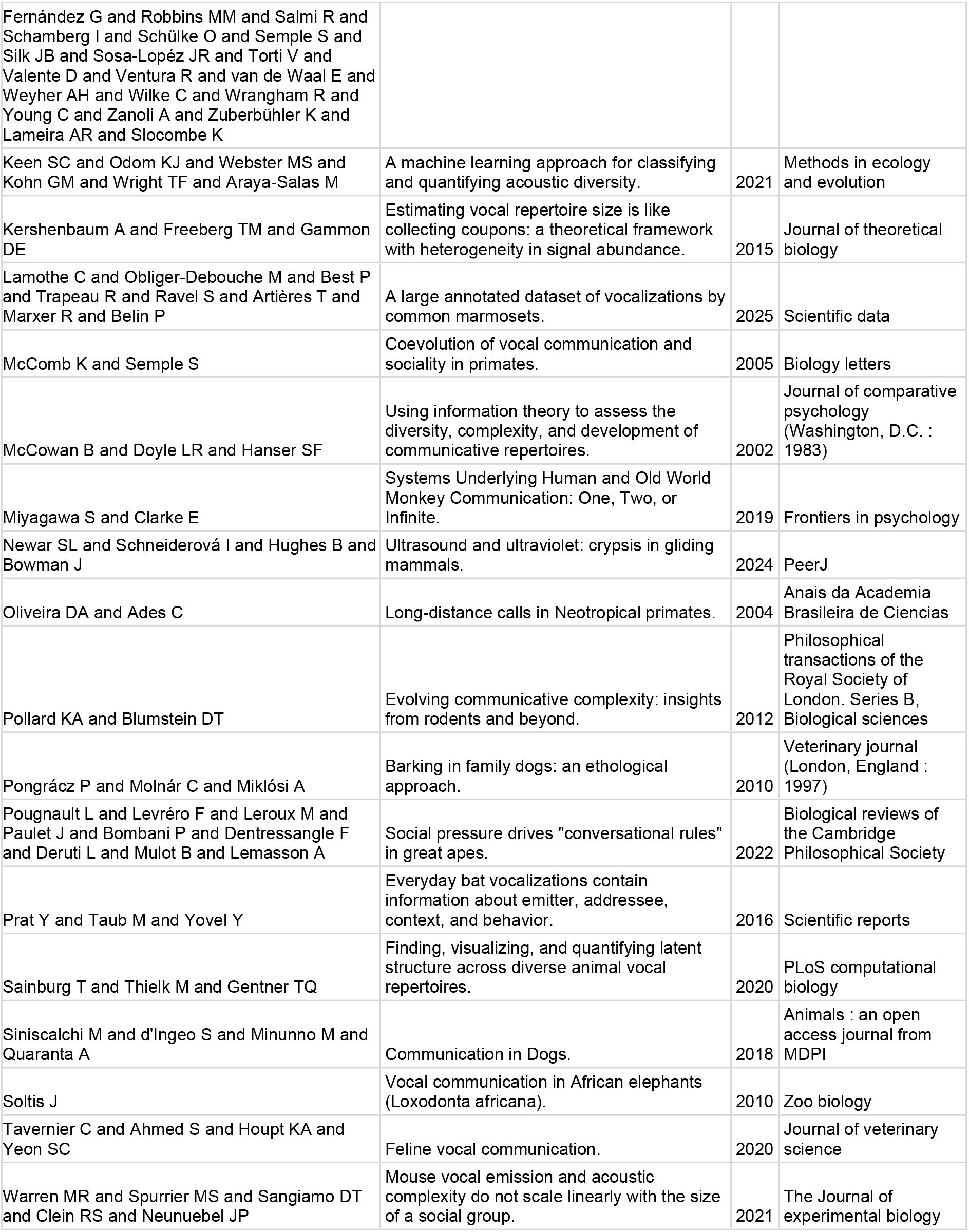

